# Quantifying spontaneous infant movements using state-space models

**DOI:** 10.1101/2024.04.16.589847

**Authors:** E. Passmore, A. K. L. Kwong, J. E. Olsen, A. L. Eeles, J. L. Y. Cheong, A. J. Spittle, G. Ball

## Abstract

Over the first few months after birth, the typical emergence of spontaneous, fidgety general movements is associated with later developmental outcomes. In contrast, the absence of fidgety movements is a core feature of several neurodevelopmental and cognitive disorders. Currently, manual assessment of early infant movement patterns is time consuming and labour intensive, limiting its wider use. Recent advances in computer vision and deep learning have led to the emergence of pose estimation techniques, computational methods designed to locate and track body points from video without specialised equipment or markers, for movement tracking. In this study, we use automated markerless tracking of infant body parts to build statistical models of early movements. Using autoregressive, state-space models we demonstrate that infant movement can be modelled as a sequence of motor states, each characterised by specific body part movements, with expression that varies with age and differs in infants at high-risk of poor neurodevelopmental outcome.

## Introduction

The emergence of motor skills in infants enriches their interaction with the environment.^1^ From lifting their head while prone, to sitting, standing and walking, motor development instigates new skills and opportunities for interaction with a child’s environment and carers, facilitating goal-directed actions and learning.^2,3^

Early motor development has cascading effects on executive function.^3^ Evidence of goal-directed movement has been observed in the developing fetus from as early as mid-gestation.^4^ From birth, as movements transition from spontaneous general movements to planned motor actions, connections are reinforced between primary motor systems and high-order cortex,^5^ accompanied by changes in functional activity of the cerebral cortex ^6^ and a shift in metabolic activity from primary sensory and motor cortex towards parietal and frontal regions.^7,8^

In school-age children, motor skills are associated with higher-order cognitive functions and academic achievement.^9–11^ Motor trajectories vary significantly across individual infants, however, and longitudinal studies have demonstrated that early postnatal motor control and earlier attainment of developmental milestones in infancy are associated with improved behavioural and educational outcomes.^2,12–14^ In contrast, disrupted motor development, in the form of abnormal postnatal general movements, is a core feature of several neurodevelopmental and cognitive disorders^15–17^ and a consequence of disruptions to early development, including preterm birth.^18^ The importance of early motor development in supporting later cognitive functions is highlighted by the promising efficacy of motor interventions in improving outcomes in at-risk populations.^19,20^

From studies of voluntary movements in humans and primates, it has been posited that movements are couched in a series of motor ‘primitives’, short kinematic elements that can be strung together and combined to form more complex actions.^21–31^ In neonates, motor primitives that are present at birth are mirrored in the basic locomotor elements of walking in both toddlers and adults.^32^ Over the first few months of life, the emergence of spontaneous, general movements: trunk rotations, and coordinated movements of the arms and legs, follow a well-defined trajectory.^33^

Between nine and twenty weeks of age, spontaneous movements are characterised by continuous ‘fidgety’ movements of the arms, legs, neck and trunk with moderate speed and variable acceleration.^34,35^ Absence or abnormality of these developmental motor patterns are a core indicator of the General Motor Assessment (GMA), an early screening tool with high predictive validity for neurodevelopmental outcomes and excellent inter-rater reliability.^34,36,37^ While the complete absence of typical fidgety movements is a good predictor of severe neurological deficits such as cerebral palsy, abnormal movements exaggerated in amplitude, speed, and jerkiness or limited in repertoire are associated with minor neurological dysfunctions and delayed motor development.^36,38,39^ Abnormal or absent fidgety movements in infancy may also be associated with delayed cognitive and language development and have been observed in genetic neurodevelopmental disorders.^39–42^

Typically, the quality of general movements in infancy can be rated using a trained assessor’s Gestalt perception from observing a supine infant, either in person or via video recording, with no direct handling or interaction.^34,36,43^ The most valid and reliable method being Prechtl’s assessment of General Movements.^37^ By contrast, quantitative kinematic analysis of body movements often rely upon specialised technology: 3D cameras, motion capture markers, or sensors.^44–46^ This can be burdensome for study participants and families and may preclude multiple, repeated assessments over time. As such, studies of motor development are often limited to relatively coarse measures of motor development (e.g.: milestone attainment).^47,48^ The development of specialised smartphone apps has enabled at-home, video-based movement assessments.^43,49^ By facilitating remote assessment, these approaches improve accessibility to clinical expertise and allow identification of high-risk infants outside of clinical settings.^36,43,49–51^

Recent advances in computer vision and deep learning have resulted in the emergence of pose estimation techniques for movement tracking, computational methods designed to locate and track body points from video, without specialised equipment or markers.^52,53^ Pose estimation tools have proven to be highly accurate and can be applied widely to different video acquisitions and tasks, including tracking infant movements.^50,54–57^ Such models require fine-tuning, or training, on infant data to accommodate significant differences in body segment size and scale compared to adults^50,56^ however, once trained, we and others have achieved promising results in predicting early motor outcomes from automated pose estimation tools.^50,54,55,58^

A common criticism of machine learning approaches in clinical settings is the opacity and limited interpretability of automated model predictions.^59^ While attention-based methods are useful to identify important periods or sequences in timeseries data they are divorced from movement mechanics.^50,55^ To aid interpretation, alternative statistical models, grounded in ethological observations that spontaneous movement can be organised as a set of movement ‘syllables’ or short motor sequences, may hold promise in the study of early human movement.^60–62^ These unsupervised, data-driven state-space models frame complex movements as a set of short sequences with unique dynamics across body points that can vary in timing and frequency across individuals, allowing the identification of motor sequences that change with age, or are affected by neurological or developmental disorders.^61,63,64^

In this study, we use automated markerless tracking of infant movements to build statistical models of early motor development. Using autoregressive, state-space models we test the hypothesis that early movement patterns can be modelled as a progression through a series of discrete dynamic states, the expression of which changes with age and is altered in infants at high-risk of poor neurodevelopmental outcomes.

## Results

### Autoregressive state-space models capture infant movement patterns

In a set of n=486 smartphone videos of supine infants (n=330 individuals, aged 12 to 18 weeks’ corrected age; n=151 born preterm; 49.4% female), we used a pre-trained deep learning model to automatically label and extract the position of 18 body points^50,65^ (**Figure 1a**). In every video frame (n=4500), each body point was represented by an *x* and *y* coordinate (normalised to infant position and size). After filtering and downsampling to 10Hz, we used Principal Component Analysis (PCA) to generate a compact representation of body movement, combining body point data into a set of Principal Movement (PMs).^66^ Each PM captures a coordinated pattern of movement across body points, reducing the dimensions of the data and accounting for redundancy between correlated body points (**Figure 1b; Figure S1)**. Overall, 15 PMs explained 89% of variance in pose (**Figure 1b**). For each video, body position in each frame was represented by a set of 15 weights (one per PM).

**Figure 1:**
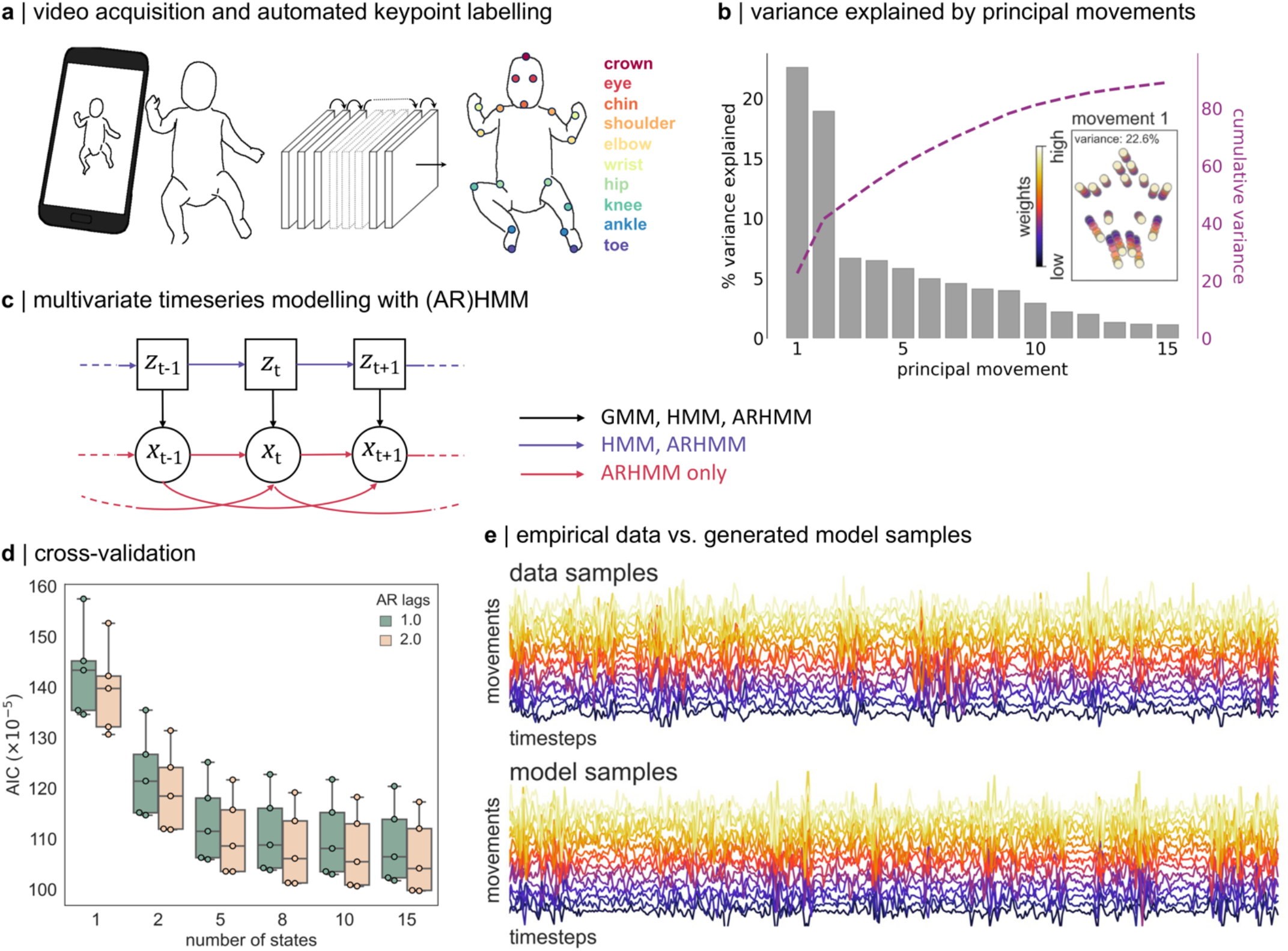
State-space modelling of infant movement dynamics. (**a**) Movement videos (3 minute length) of infants aged between 12 and 18 weeks of age were acquired at home using a specialised smartphone app (Baby Moves). Using a custom-trained deep learning algorithm (DeepLabCut), video frames were automatically labelled to track several key body points. (**b**) Following preprocessing and quality control, movements were represented by a set of Principal Movements (PMs). Plots shows the variance explained (cumulative variance; right axis) by each PM in a random subset of n=100 videos. Inset: First PM. Principal movement is shown by position of markers at different weights. (**c**) The dynamic contribution of PMs to bodypoint movement in each video was modelled using state-space models. Graphical model shows dependence of observations, *x*, on the transition between states, *z*, and previous values of *x* at *t-*1 and *t-*2. In HMM, autoregressive components are removed. In GMM, state progression is removed. (**d**) Goodness-of-fit was compared between models using 5-fold cross-validation. Plots shows AIC across folds for ARHMM models with different values of lag (lower is better). (**e**) the first 500 frames of a randomly selected video are shown (top). Each line shows the first derivative of a given principal movement over time. Bottom: Synthetic data. 500 frames generated from the trained AR(2)HMM model (k=8) show observed movement dynamics are captured by the state-space model.

Using the first derivative of PM weight over time, we modelled changes in movement velocity using state-space models, evaluating goodness-of-fit with 5-fold cross-validation (**Figure 1c-d**; **Figure S2**). We performed initial model selection using a random subset of n=100 videos (ensuring a maximum of one video per participant was selected) as an evaluation set, comparing model performance between hidden Markov models with and without autoregressive terms (ARHMM and HMM, respectively) and between simple Gaussian models without state progression (GMM) (**Figure 1c**). These data-driven approaches for clustering multivariate timeseries identify periods with similar movement dynamics across videos.

For each model class, we evaluated model fit over a number of states from k=1 (i.e.: all movement is generated from a single set of autocorrelation dynamics) up to k=15. For all models, performance in held-out test data (averaged over cross-validation folds) improved when including additional states, with little significant improvement above 5 to 8 states (**Figure S2; Table S1**). The number of states indicates the timescale of observed temporal structure within the movement data and provides a natural segmentation of continuous behaviour into meaningful components of movement.^61^

Overall, model performance was higher using ARHMM models compared to HMM and was, in turn, higher than in GMM models (**Figure S2**). Under the simpler GMM model, infant movement dynamics are modelled as variations of a set of poses or positions with no additional temporal structure. HMM models include additional transition probabilities, testing the hypothesis that behavioural modules, or movements, transition from one to another with a specific probability that is consistent over subjects. The addition of autoregressive (AR) coefficients tests module-specific dependencies of movement along a shorter, sub-second timescales. The improved performance of ARHMM models supports the hypothesis that spontaneous infant movements can be described with a hierarchical process, with movements grouped into modules encoding the occurrence of stereotyped behaviours or movement patterns across individuals, the dynamics of which are governed by autocorrelation over a short timescale. Increasing the autoregressive order improved ARHMM model performance moderately (**Figure 1d**). Based on average performance over folds in the evaluation set, the best fit model was selected as an 8-state ARHMM with lag=2. This model was then fit on the full dataset for further analysis. We confirmed that this model was able to generate synthetic data samples with feature distributions and dynamics matched to empirical data (**Figure 1e; Figure S3**).

### The expression of motor states in early infancy

Using the fitted model, we estimated the framewise state-membership for each video (**Figure 2a**), identifying periods where movements followed a given state dynamic. On average, state transitions were rapid with each state capturing only small segments of each video. Across all videos, the median state length (excluding single frame instances) was 3 frames (0.3sec; **Figure 2a**) with the average [IQR] maximum time spent in any given motor state = 19 [15.75-26.0] frames (1.9sec). State occupancy (proportion of time spent in each state) varied across individual videos (**Figure 2a-b**).

**Figure 2:**
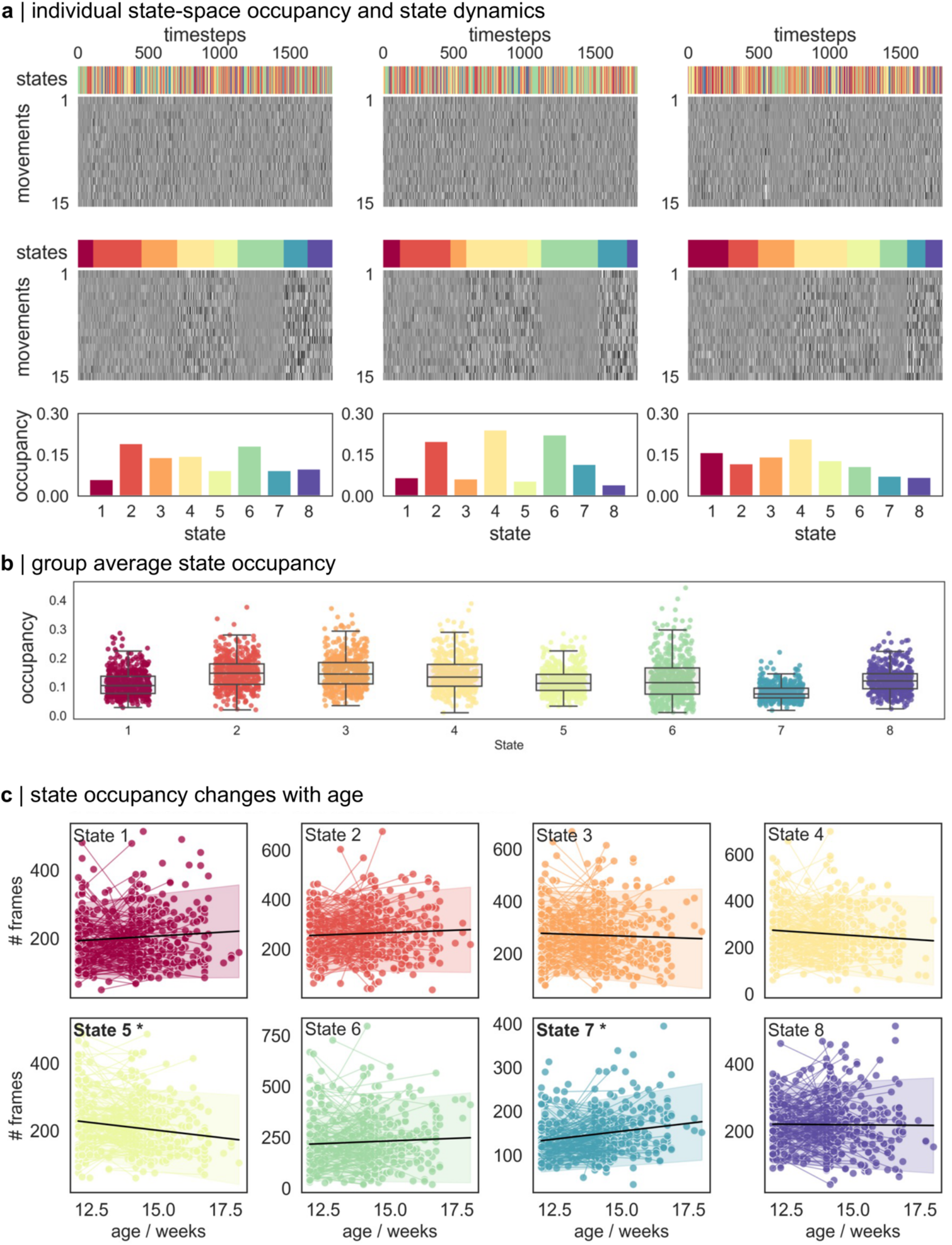
Changes in state occupancy during infant movement. (**a**) Individual movement dynamics (n=3, one video per column) are modelled using best-fit ARHMM. Top row: most likely states for each video frame. Movement dynamics are shown below in grayscale. Each row represents weight of a given PM in a given frame. Middle row: Frames are sorted according to state membership. Different movement dynamics are apparent across states. Bottom row. State occupancy (% frames in each state) for each video. (**b**) Group average state occupancy. For each video (n=486) average state occupancy is shown. Boxplot indicate median and interquartile range. (**c**) Change in state-occupancy with age was modelled using linear mixed effects models. Videos from the same individual are joined by a line. Black line indicate main effect of age with 95% confidence intervals shaded. Significant effects (p<0.05 after Bonferroni correction over 8 states) are highlighted in bold.

Using linear mixed effect models, we tested whether the expression of specific movement states changed with age. We found that between 12 and 18 weeks of age, the number of state 5 movements decreased (*β*_age_= -9.26; p <0.001), whereas the number of state 7 movements increased (*β*_age_= 7.23, p<0.001; **Figure 2c**; **Table S2**). Small increases were also observed in states 1 (p=0.03) and 4 (p=0.01), though these did not pass correction for multiple comparisons.

### Early movement patterns differ in high-risk infants

Participant’s motor development was assessed using the GMA (n=326/330) by two independent trained assessors. We next tested if state occupancy varied between participants with normal (n=289) compared to abnormal (sporadic, absent or abnormal, n=37) fidgety general movements. Where two videos were available, the GMA was based on the video acquired at the later timepoint.^35,50^ As preterm birth is an independent risk factor for poor motor development, we included birth status (preterm normal/abnormal = 117/31; term control normal/abnormal = 172/6) along with age in the model as additional factors.

Movements in state 7 were significantly increased in participants with abnormal or absent GMs (*β*_GM_ = 23.7, p=0.003; **Table S3**. Preterm birth was associated with increased occupancy of states 4 and 5 (*β*_birth_ = 38.7, 26.8, p <0.001, =0.001, respectively) and a decreased count of state 1, 3 and 8 movements (*β*_birth_ = -31.9, -35.8, -34.4, respectively, p <0.001; **Table S3**).

At 2 years of age, follow-up assessments were available for 302/330 (motor and cognitive), 272/330 (language) participants. We found that lower motor scores at 2 years were associated with increased occupancy of state 7 in infancy after adjusting for preterm birth status, although this association did not pass correction for multiple comparisons (*β*_motor_ = -0.39, p =0.014; **Table S4**). Improved cognitive scores and language scores were associated with lower occupancy of state 4 (*β*_cog_ = -1.19, *β*_lang_ -0.85, p =0.0012, 0.0051, respectively) and state 1 (*β*_lang_ -0.67, p =0.003; **Table S5-6**).

### Specific motor states capture high velocity body movements

To examine movement dynamics of each motor state, we identified all frames in each state for every participant video. A random sample of state-specific movements (n=5000, frame-to-frame movement of each body point) is shown in **Figure 3a**. The magnitude of bodypoint movements in each state vary significantly, with several states capturing high velocity (large magnitude) movements between frames (e.g.: states 4, 7 and 8), and other signifying periods of low motion, or rest (states 2, 3 and 6). State 7 captures high velocity movements of bodypoints in all directions, including large magnitude movements of feet (ankles, toes) and hands (wrist).

**Figure 3:**
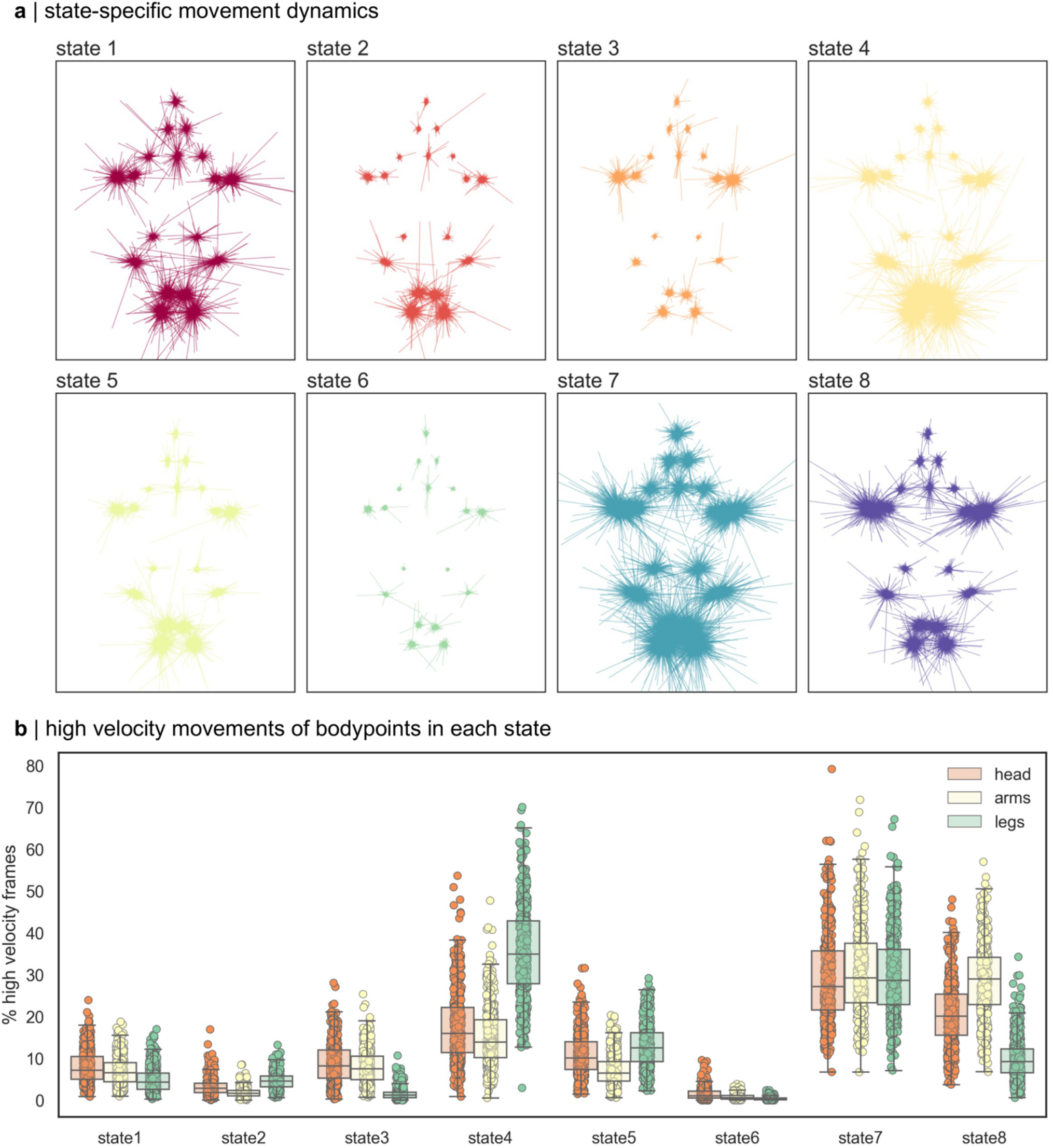
State movement dynamics. (**a**) Direction and magnitude of frame-to-frame movements of each body point in each state, centered on the group average position. Movements were randomly selected (n=5000) from frames in each state in all videos. (**b**). For each body point, the top 10% high velocity frames (those containing most movement) were identified and grouped by state membership. Plots shows average % high velocity frames for each group of bodypoints in each state for every video (n=486). Boxplot shows median and interquartile range.

To identify potential specificity of movements across bodypoints in each state, we identified the top 10% of frames and corresponding states with the largest frame-to-frame motion in each bodypoint, grouped into three larger clusters: arms (shoulder, elbow, wrist), legs (hips, knee, ankle, toe) and head (crown, chin, eyes) (**Figure 3b**). Of the three high-motion states (4,7 and 8), we found that each state captured different patterns of movement, with states 4 and 8 preferentially capturing high velocity leg and arm movements, respectively, in contrast to the whole-body movements of state 7. State 3 and 5 were associated with smaller magnitude movements of the arms and legs, respectively. States 2 and 6 largely signified periods of low movement, capturing few high velocity frames across body points.

## Discussion

In this study, we combined deep learning-based video tracking with autoregressive state-space models to identify dynamic movement states in infants aged 12 to 18 weeks. We observed that infant movement patterns can be modelled as a sequence of motor states, each characterised by specific body part movements, with expression that varies with age and differs in infants at high-risk of poor neurodevelopmental outcome.

Early motor development has been extensively studied, with the timing and trajectory of major movement milestones defined across ages and populations.^48^ In the months following birth, several characteristic movement periods have been defined, the occurrence and calibre of which offer prognostic value for later developmental and neurological outcomes.^16,33–35,39,40,42^ Alongside others, we have recently demonstrated that automated methods, founded upon recent computer vision and machine learning innovations,^52,65^ are able to capture body part movements and quantify risk in infant populations using video alone, without the need for specialised tracking equipment.^50,54,58^ While powerful, a current limitation of these methods is the difficulty interpreting algorithmic model decisions. Here, we combined AI-based video tracking with a statistical modelling framework previously used in ethological studies of spontaneous animal movement.^60,62–64,64^ These data-driven methods are able to parse complex, multivariate movement traces, identifying shared patterns of motor behaviour that are common to study participants. In turn, this allows the analysis of timing, frequency and progression of each motor state over time and the quantification of individual variations in movement.^61,63,64^

Hidden Markov Models describe a system where observable, external events depend upon a set of internal, unobservable factors. In the context of this study, unable to observe the internal factors that lead to different motor behaviours in the infant, we infer the identity of a series of modules, or states, that occur in sequence and are each associated with a specific type of movement observable in the video data. We find that HMM models outperform simpler GMM models, that do not account for the sequential progression of internal states and that the addition of autoregression terms, to account for the correlation in movements between adjacent frames, confers a significant benefit. Once fitted to the infant movement data, we find that ARHMMs can produce synthetic multivariate timeseries data that share distributional and dynamic features with the empirical data (**Figure 1e**). Our ARHMM models demonstrate that a relatively small number of states (between 5 and 8) are sufficient to explain the full complement of infant movements, outperforming simple models based on covariance and autocorrelation between body parts alone (i.e.: k=1 state), or more complex models with higher number of internal states (k=10, 15).

The notion that complex motor behaviours can be modelled as a combination of simpler movement ‘syllables’ or ‘primitives’ is long-standing.^25,27,28,32,32,61^ The states defined in this study can be viewed in a similar way: as compositional elements of complex movements. By segmenting multivariate movement traces into a set of modules, we can isolate particular movement patterns, each governed by its own set of autocorrelation dynamics, that reflect a stereotyped body movement – fast, slow, specific or global. In this study, we describe eight motor states, or coordinated movement patterns, three of which are associated with high velocity, or large magnitude, movements between adjacent frames. Others are associated with small movements, or pauses in movement, or capture high velocity movements in specific body parts, i.e.: arms or legs (**Figure 3**). While some states captured up to 40-50% of frames in a single video, the median time spent in a single state was very short with the maximum time spent before switching around 2 sec on average across individuals. The frequent transitions between distinct movements provide a granular description of motor behaviours with body part position dictated by the interaction of specific movement types and the location of body points in the preceding frames. These movement states, when strung together, encode the full repertoire of transient, spontaneous infant movements.

We found that large, high velocity, whole-body movements (state 7) increased in frequency with age. This movement type was also associated with neurodevelopmental risk, increased in infants with abnormal or absent fidgety general movements and associated with lower motor scores at 2 years. In contrast, movements in state 4 and 5, involving high-amplitude movements of the legs were reduced in older infants but increased in those born preterm, suggesting a potential difference in developmental timing in high-risk infants.^35,67–69^ Abnormal fidgety movements that are exaggerated in amplitude and speed are a known risk factor for poor neurodevelopmental outcome.^40,70,71^ State 7 movements may reflect such exaggerated whole-body movements although the alignment between motor state timing and specific pathological movement types requires further validation. We lack the high resolution annotation of movements within each video needed for a direct comparison between motor state and movement type. However, others have demonstrated that automated video tracking methods are able to detect abnormal periods associated with specific types of GM, such as fidgety movements.^72^ While it is unlikely that a single motor state would fully capture specific abnormal movements observed with the GMA, our study indicates that state-space models of infant movement are able to parameterise complex, spontaneous infant movements and are sensitive to abnormalities or absence of typical developmental movements. Combination with other, extended motor assessments such as the Motor Optimality Score (revised; MOS-R), may also provide further insight into the importance of specific movement patterns, posture and quality observations during this time period.^73–75^ We propose that this approach will prove useful to the automated detection and improved understanding of atypical motor behaviour in high-risk infant populations.^50,54,58^

## Acknowledgements

We would like to acknowledge the parents/families of infants who participated in the study. We would also like to acknowledge the extended Victorian Infant Collaborative Study team for their contribution in collecting infant and 2-year follow up data. We would like to acknowledge funding from the Rebecca L Cooper Medical Research Foundation (PG2019421 to G.B.), National Health and Medical Research Council Investigator Grant (1194497 to G.B., 2016390 to J.L.Y.C.), NVIDIA Corporation Hardware Grant program, The Royal Children’s Hospital Foundation, Melbourne and the Murdoch Children’s Research Institute Clinician Scientist Fellowship.

## Author contributions

Recruitment and data acquisition: A.L.K., J.E.O., A.L.E., J.L.Y.C., A.J.S. Data processing: A.L.K., E.P., G.B. Data analysis: E.P., G.B. Resources: E.P. G.B. J.L.Y.C., A.J.S. Supervision: A.J.S., G.B. Writing: E.P, G.B. Revising and editing: all authors

## Code and data availability

Analysis code supporting this study is available at https://github.com/garedaba/state-space. Requests for data that supports the findings of this study can be made to the Murdoch Children’s Research Institute Data Office. The data is not publicly available due to privacy and ethical restrictions.

## Materials & Methods

### Participant information

In total, we acquired video data from parent-participants from the Victorian Infant Collaborative Study (2016/2017 cohort) n=341 infants aged between 12- and 18-weeks’ corrected age. This included n=155 infants born extremely preterm (< 28 weeks’ gestation; 77 [50%] female; mean age ± SD = 26.8 ± 2.0 weeks) and n=186 term-born control infants (37-42 weeks’ gestation; (91 [49%] female; mean age ± SD = 39.5 ± 1.2 weeks). In 165 infants (78 preterm; 87 term), two videos were acquired, resulting in a total of 506 videos. Following initial QC after automated labelling, we excluded 18 videos (see **Trajectory data processing** below) and excluded a further two videos due to missing age data (1) and a high video resolution that precluded automated labelling, resulting in a final dataset of n=486 videos acquired from 330 infants (151 preterm; 163 female). Full details of the study protocol can be found in Spittle et al.^43^ The study was approved by the Royal Children’s Hospital Ethics Committee (HREC35237).

### Video acquisition

Infant movement videos were captured using a dedicated smartphone app, Baby Moves, by a parent/caregiver at home on their personal device between April 2016 and May 2017, as detailed previously.^43,50^ Briefly, following in-app guidance, infants were recorded lying quietly in a supine position for 3 minutes. Videos were uploaded to a secure database in MP4 format. Due to differences in device model and settings, videos were acquired at different resolutions with a median frame rate of 30 (range: 15-31) frames per second. Each video was 3 minutes in length resulting in mean ± S.D frames per video of 5100 ± 497.^50^

### Markerless movement tracking

We implemented a custom keypoint labelling system based on DeepLabCut (v2.1)^65^ to track infant body part movements from video, for details see: Passmore et al.^50^ In brief, a deep learning model was trained to identify 18 body points (head, eyes, chin, shoulders, elbows, wrists, hips, knees, heels and halluces) in each video frame. For each frame, the predicted *x*, y pixel coordinates of each point are returned along with a measure of prediction confidence.^65^ Points with a confidence <0.2 were removed on a framewise basis.^50^

We have previously shown that this system results in human-level labelling accuracy, with a mean error in unseen data of 6.8 pixels (human inter-rater error = 6.9 pixels) and is robust to different video resolutions and frame rates, as well as infant clothing, background, and lighting.^50^

### Trajectory data processing

Following automated labelling, keypoint data were processed following a custom pipeline consisting of quality control (QC), outlier removal, gap filling, adjustment for camera movement and scaling.^50^ For initial QC, videos with less than 70% of keypoints labelled on average across frames were removed, resulting in n=18 excluded videos. Within each video, keypoint outliers were identified using a two-step process, first identifying labels that lay outside of an ellipse centered and scaled to infant position and size, then identifying labels that lay outside of body part-specific ellipses centred on each keypoint’s median position over time.^50^

After QC, gaps due to missing, excluded or occluded labels were filled using linear interpolation (for gaps of < 5 frames), or an iterative multivariate imputation.^76,77^ As videos were acquired from handheld devices, we accounted for camera movement by applying a rotation to all points to align the hips and shoulders across frames . To account for differences in infant size, keypoint coordinates were scaled to infant length (defined as distance from crown to mid-hip).

For each frame, we extracted the normalised *x* and *y* coordinate of each keypoint, resulting in *d* = 36 pose features per frame. The timeseries each feature was bandpass filtered using a fourth-order, zero-phase Butterworth filter (0.01 - 5 Hz) to reduce low-frequency-drift and the high-frequency jitter commonly associated with automated framewise labelling. After removing drift, timepoints where *x* or *y* position of a given keypoint were > 3 standard deviations from the median position of the whole cohort were identified as outliers and, to reduce computational burden, the length of each timeseries downsampled to 1800 frames (10Hz frame rate) using a first-order spline interpolation, excluding any outlying points.

### Principal movements

Pose features data from each video can be represented as a *t* timepoint × d feature (1800 × 36) matrix, *M*. As body part movements are highly correlated over time, we can apply PCA to generate a more compact representation of movement by a smaller number of basis components that each represent a coordinated pattern of movement across body parts, or Principal Movements (PM).^66^

To estimate a set of PMs, we randomly selected n=100 videos (ensuring only one video was selected for any given participant). Each feature timeseries was demeaned to remove body position and normalised to unit standard deviation to equalise the contribution of each feature to the decomposition. Normalised coordinate data were concatenated along the time dimension and decomposed with SVD:

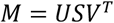

Resulting in a set of pose eigenvectors, or PMs, *V* = {*v*_1_, …, *v*_*k*_}, and weights, *US* = {*ε*_1_, …, *ε*_*k*_} that, for a given mode *k*, quantify the degree to which posture at time *t* deviates from the mean pose in the direction of *v*_*k*_. Unseen data can be projected onto this basis set through multiplication with the set of calculated eigenvectors. Principal movements are shown in **Figure S1**.

### State-space modelling

Individual movement dynamics were modelled using an autoregressive Hidden Markov Model (ARHMM), a data-driven approach to multivariate timeseries clustering. Using Hidden Markov Models (HMM), a continuous, *d*-feature × *t* timeseries, *X*, can be modelled as the output of a Markov process, *Z*, with a set of *K* hidden states, {*z*_1_, *z*_2_, …, *z*_*k*_}, that are not directly observable. Each state is associated with a *d*-dimensional mean, *μ*_*k*_, and a *d* × d covariance matrix, *S*_*k*_ from which the multivariate observations are drawn at each time step. The hidden states are assumed to progress as a Markov process where the next state is only dependent on the current state: *p*(*z*_*t*_|*z*_*t*−1_). Switches between states are governed by a *K* × *K* transition matrix, *T* containing state transition probabilities, *Φ*, where *Φ*_*zz*,_ indicates the probability of transitioning from state *z* to state *z*’ at the next timestep. In HMMs, the outputs, or observations, are determined solely by the given state at each time step and therefore do not capture short term correlations in timeseries data.

In an ARHMM, the observations at time *t, x*_*t*_, are dependent on both the hidden state, *z*_*k*_, and the observations at previous timepoints, {*x*_*t*−1_, x_*t*−2_, …, x_*t*−*l*_} with *l* determined by the degree of lag, *L*, specified. As such the observations for a given state, *z*_*k*_, are defined as:

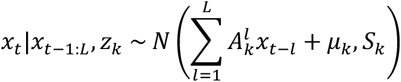

Where *x*_*t*_ is a set of observations at time *t*, 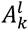 is a matrix containing the linear dynamics (the relationship between observations at *t* and at a given lag, *l*) for a given state, *μ*_*k*_ is a state-dependent bias (average value of each variable in each state) and *S*_*k*_, a state-dependent covariance function. The number of hidden states and the degree of lag considered in each state are hyperparameters for the model. To focus on dynamic movement, ARHMM was applied to the first derivative (i.e.: velocity) of each timeseries.

### Cross-validation

Model performance was initially assessed using 5-fold cross-validation across n=100 randomly selected videos. In each fold, we performed a grid search across ARHMM hyperparameters: *L* = [1, 2] and *K* = [1, 2, 5, 8, 10, 15], where *K* = 1 represents a special case of a standard *AR*(*l*) model (i.e.: no state progression). As baseline models, we also tested standard HMM models (i.e.: no autoregression) across a range of *K* as well as Gaussian Mixture Models (GMM) where observations are drawn from a set of *K* multivariate distributions (i.e.: no state progression or autocorrelation). Assessment of model performance was based on average log-likelihood of test data samples given the model, averaged over five folds. Models were fit using stochastic expectation-maximisation for a maximum of 500 iterations or until convergence. After cross-validation, the best performing model was fit on all data samples (n=486 videos; max. 500 iterations). To account for stochasticity in the model fitting procedure, final model fitting was repeated 25 times and the model parameters averaged. All (AR)HMM and GMM models were implemented using the **ssm**^78^ and **scikit-learn**^77^ packages, respectively.

### Motor assessment and follow-up

To assess motor development at the time of video acquisition, infant videos were independently assessed using Prechtl’s General Motor Assessment (GMA).^34^ Assessments were performed by two independent, trained assessors, blinded to each participant’s neonatal history, viewing each video. General movements (GMs) were classified as normal if fidgety GMs were intermittently or continuously present, absent if fidgety GMs were not observed or were sporadically present, or abnormal if fidgety GMs were exaggerated in speed and amplitude. Disagreements were resolved by a third experienced GMA trainer and assessor who made the final decision. Any videos rated as unscorable were not evaluated in this study. GMA scores were available for 326/330 infants.

Neurodevelopmental follow-up was performed at 2-years’ corrected age using the Bayley Scales of Infant and Toddler Development-3rd edition (Bayley-III) motor, cognitive and language domains. Bayley-III scores were available for 302/330 infants for motor and cognitive domains and 272/330 infants for the language domain.

### Statistical analysis

Linear mixed effects models were used to test main effects of age, birth status (preterm or term) and neurodevelopmental outcomes (GMA and Bayley-III) on state occupancy. To account for repeated measures from multiple videos acquired in the same subject, participant ID was included as a random effect. For each analysis, statistical significance was considered at p<0.05 after Bonferroni correction for multiple comparison over states (i.e.: 0.05 / 8).

Analyses and visualisations were implemented in **Python (3.10)** using packages including **numpy**^79^, **scipy**^80^, **scikit-learn**^77^, **ssm**^78^, **matplotlib**^81^, **seaborn**^82^, **pandas**^83^ and **einops**.^84^

