## Supplemental Information for "Quantifying spontaneous infant movements using state-space models"

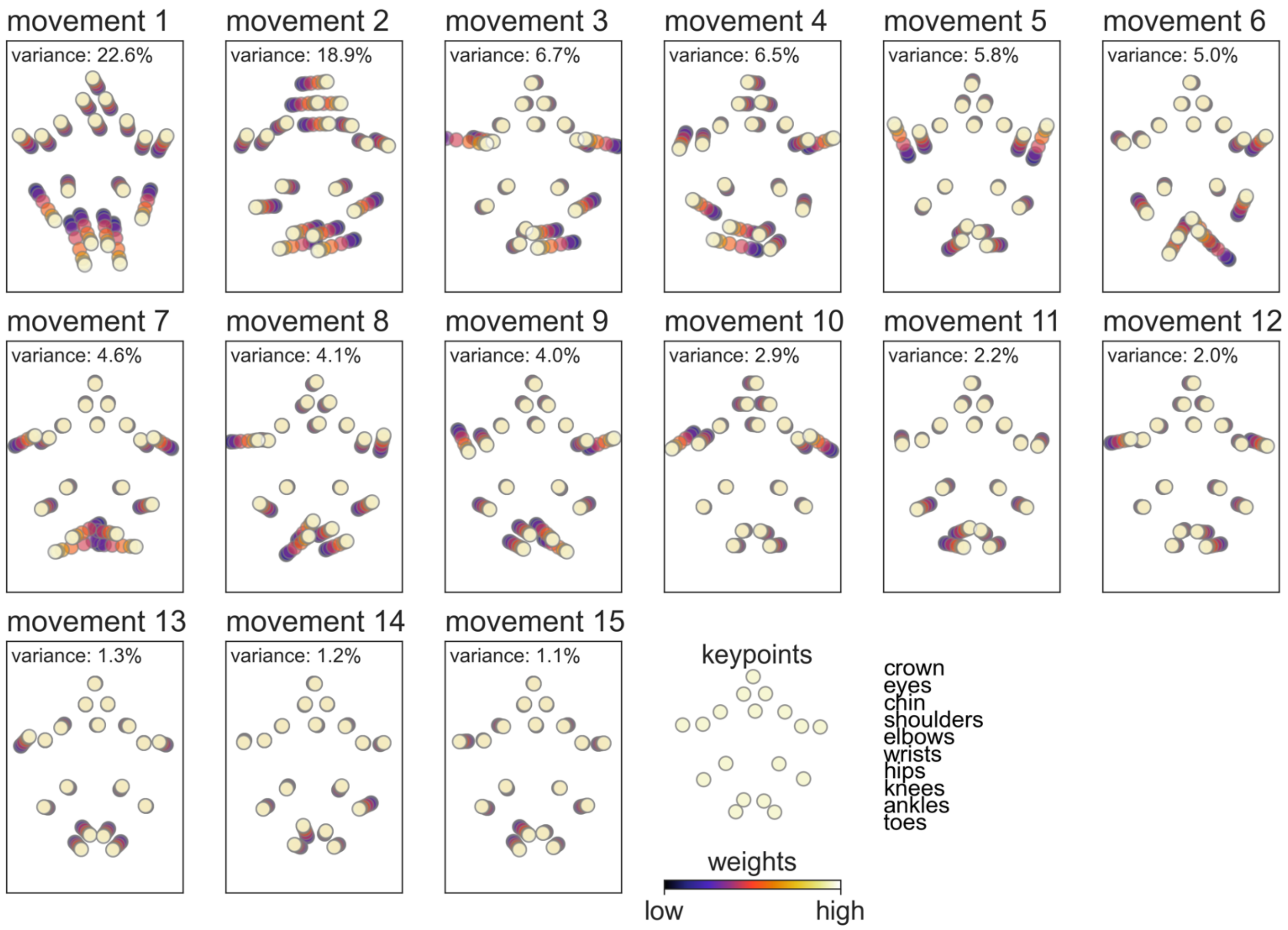


**Figure S1: Principal movements.** Each plot shows one principal movement (PM). The position of each keypoint at a given PM weight is shown in colour (colourbar, arbitrary scale).

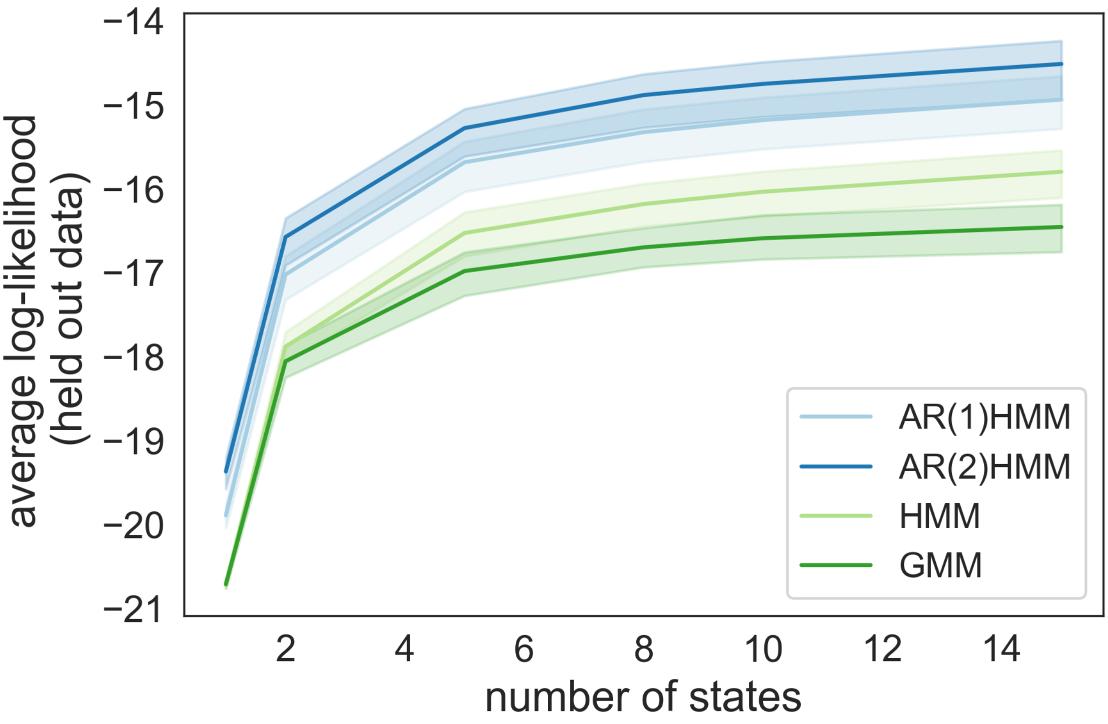


**Figure S2: Goodness-of-fit for HMM and GMM models.** Average log-likelihood on the test set across 5 cross-validation folds is shown with 95% confidence intervals (shaded) for each model type.

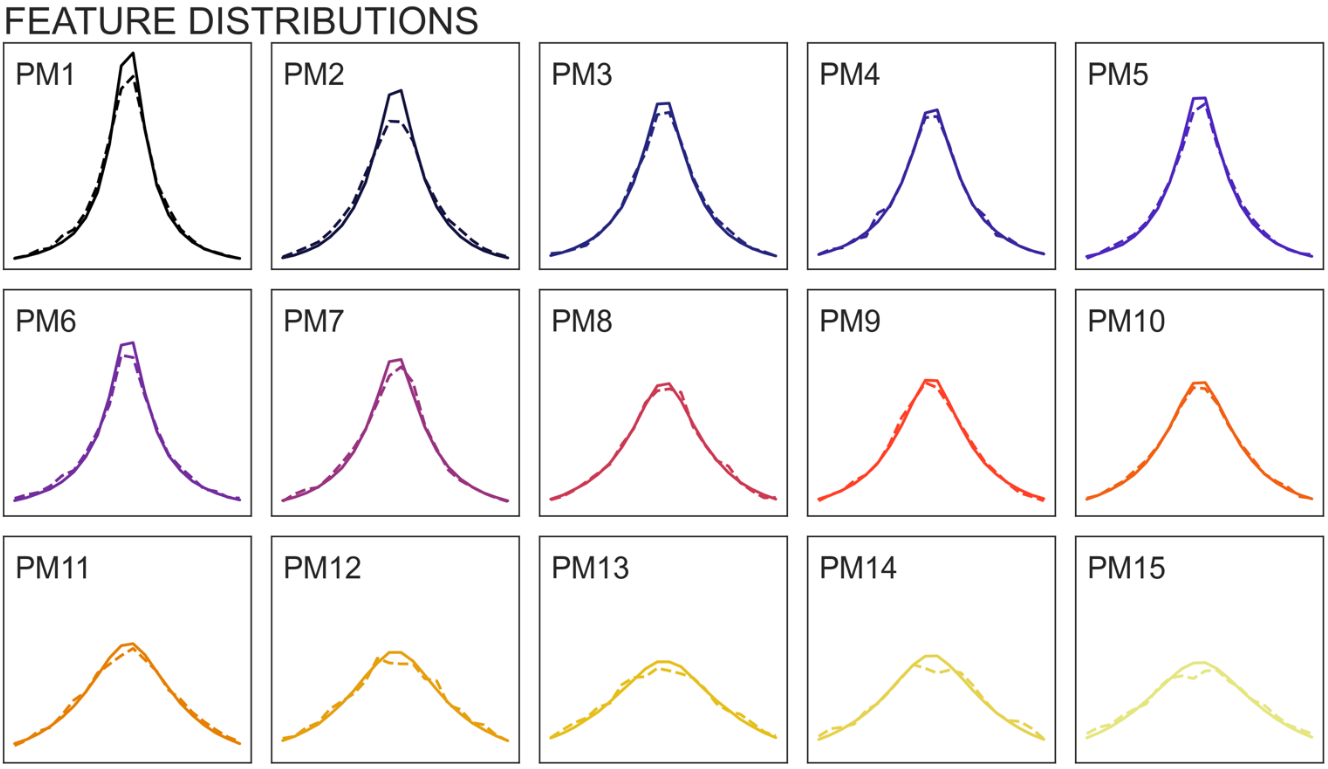


**Figure S3: Distribution of PM weights in empirical (solid) and synthetic (dashed) data.**

**Table S1: model fit averaged across cross-validation folds**

| model | states | lag | log-likelihood | AIC |
|---|---|---|---|---|
| GMM | 1 | - | -20.71 | 1490884 |
| | 2 | - | -18.05 | 1300229 |
| | 5 | - | -16.98 | 1223700 |
| | 8 | - | -16.70 | 1204180 |
| | 10 | - | -16.59 | 1196997 |
| | 15 | - | -16.45 | 1188694 |
| HMM | 1 | - | -20.71 | 1491098 |
| | 2 | - | -17.88 | 1288119 |
| | 5 | - | -16.53 | 1192168 |
| | 8 | - | -16.18 | 1168899 |
| | 10 | - | -16.04 | 1159374 |
| | 15 | - | -15.80 | 1144988 |
| AR(1)HMM | 1 | 1 | -19.89 | 1432452 |
| | 2 | 1 | -17.02 | 1227143 |
| | 5 | 1 | -15.68 | 1133670 |
| | 8 | 1 | -15.33 | 1110840 |
| | 10 | 1 | -15.18 | 1102360 |
| | 15 | 1 | -14.94 | 1089789 |
| AR(2)HMM | 1 | 2 | -19.37 | 1395196 |
| | 2 | 2 | -16.57 | 1195808 |
| | 5 | 2 | -15.28 | 1106586 |
| | 8 | 2 | -14.88 | 1082540 |
| | 10 | 2 | -14.75 | 1075781 |
| | 15 | 2 | -14.51 | 1065919 |

**Table S2: Fixed effects model terms for age-related changes in state occupancy**

| model terms | intercept | | age | |
|---|---|---|---|---|
| state | β | p | β | p |
| 1 | 137.62 | $6.47\times10^{-6}$ | 4.65 | $3.10\times10^{-2}$ |
| 2 | 208.04 | $7.35\times10^{-8}$ | 3.91 | $1.53\times10^{-1}$ |
| 3 | 318.90 | $8.58\times10^{-14}$ | -3.43 | $2.57\times10^{-1}$ |
| 4 | 364.26 | $1.48\times10^{-17}$ | -7.55 | $1.26\times10^{-2}$ |
| 5 | 339.41 | $5.01\times10^{-30}$ | -9.26 | $1.08\times10^{-5}$ |
| 6 | 153.50 | $1.95\times10^{-3}$ | 5.27 | $1.33\times10^{-1}$ |
| 7 | 46.11 | $1.91\times10^{-2}$ | 7.23 | $2.02\times10^{-7}$ |
| 8 | 229.15 | $4.80\times10^{-13}$ | -0.66 | $7.68\times10^{-1}$ |

**Table S3: Fixed effects model terms for associations with GMA outcome**

| model terms | intercept | | age | | birth status | | GMA outcome | |
|---|---|---|---|---|---|---|---|---|
| state | β | p | β | p | β | p | β | p |
| 1 | 124.66 | $1.22\times10^{-4}$ | 4.58 | $3.65\times10^{-2}$ | -31.85 | $1.11\times10^{-4}$ | 3.00 | $8.14\times10^{-1}$ |
| 2 | 212.34 | $2.21\times10^{-7}$ | 4.32 | $1.22\times10^{-1}$ | 21.52 | $2.86\times10^{-2}$ | -2.41 | $8.74\times10^{-1}$ |
| 3 | 288.88 | $1.16\times10^{-10}$ | -4.89 | $1.10\times10^{-1}$ | -35.81 | $7.81\times10^{-4}$ | -34.00 | $3.87\times10^{-2}$ |
| 4 | 395.04 | $1.52\times10^{-18}$ | -6.15 | $4.49\times10^{-2}$ | 38.69 | $2.99\times10^{-4}$ | 32.77 | $4.72\times10^{-2}$ |
| 5 | 340.24 | $7.25\times10^{-27}$ | -8.72 | $4.21\times10^{-5}$ | 26.82 | $1.09\times10^{-3}$ | -7.83 | $5.38\times10^{-1}$ |
| 6 | 150.43 | $4.38\times10^{-3}$ | 4.73 | $1.82\times10^{-1}$ | 22.43 | $1.04\times10^{-1}$ | -23.31 | $2.76\times10^{-1}$ |
| 7 | 61.40 | $3.47\times10^{-3}$ | 7.33 | $2.65\times10^{-7}$ | -7.80 | $1.26\times10^{-1}$ | 23.72 | $2.64\times10^{-3}$ |
| 8 | 222.96 | $2.60\times10^{-11}$ | -0.98 | $6.68\times10^{-1}$ | -34.38 | $1.65\times10^{-5}$ | 8.45 | $4.93\times10^{-1}$ |

**Table S4: Fixed effects model terms for associations with Bayley Motor scores**

| model terms | intercept | | age | | birth status | | Motor | |
|---|---|---|---|---|---|---|---|---|
| state | β | p | β | p | β | p | β | p |
| 1 | 178.29 | $1.23\times10^{-5}$ | 2.83 | $1.95\times10^{-1}$ | -31.66 | $1.80\times10^{-4}$ | -0.32 | $2.07\times10^{-1}$ |
| 2 | 175.73 | $6.26\times10^{-4}$ | 4.31 | $1.25\times10^{-1}$ | 20.48 | $4.65\times10^{-2}$ | 0.36 | $2.39\times10^{-1}$ |
| 3 | 223.25 | $8.42\times10^{-5}$ | -1.86 | $5.50\times10^{-1}$ | -30.29 | $7.70\times10^{-3}$ | 0.54 | $1.10\times10^{-1}$ |
| 4 | 469.51 | $2.77\times10^{-16}$ | -8.68 | $5.38\times10^{-3}$ | 32.68 | $4.74\times10^{-3}$ | -0.67 | $5.09\times10^{-2}$ |
| 5 | 333.96 | $6.63\times10^{-16}$ | -8.64 | $6.47\times10^{-5}$ | 24.72 | $4.94\times10^{-3}$ | 0.11 | $6.65\times10^{-1}$ |
| 6 | 138.09 | $4.61\times10^{-2}$ | 6.87 | $5.74\times10^{-2}$ | 20.57 | $1.65\times10^{-1}$ | 0.03 | $9.43\times10^{-1}$ |
| 7 | 93.19 | $3.24\times10^{-4}$ | 6.34 | $5.51\times10^{-6}$ | -9.11 | $8.33\times10^{-2}$ | -0.39 | $1.39\times10^{-2}$ |
| 8 | 185.48 | $1.05\times10^{-5}$ | -1.19 | $6.05\times10^{-1}$ | -27.59 | $1.08\times10^{-3}$ | 0.35 | $1.69\times10^{-1}$ |

**Table S5: Fixed effects model terms for associations with Bayley Cognitive scores**

| model terms | intercept | | age | | birth status | | Cognitive | |
|---|---|---|---|---|---|---|---|---|
| state | β | p | β | p | β | p | β | p |
| 1 | 206.30 | $7.61\times10^{-7}$ | 2.74 | $2.09\times10^{-1}$ | -33.80 | $4.61\times10^{-5}$ | -0.59 | $2.88\times10^{-2}$ |
| 2 | 178.04 | $7.25\times10^{-4}$ | 4.27 | $1.28\times10^{-1}$ | 19.61 | $5.38\times10^{-2}$ | 0.35 | $2.97\times10^{-1}$ |
| 3 | 197.08 | $6.94\times10^{-4}$ | -1.78 | $5.66\times10^{-1}$ | -28.84 | $9.73\times10^{-3}$ | 0.80 | $2.85\times10^{-2}$ |
| 4 | 521.72 | $3.82\times10^{-19}$ | -8.80 | $4.64\times10^{-3}$ | 28.92 | $1.04\times10^{-2}$ | -1.19 | $1.24\times10^{-3}$ |
| 5 | 355.22 | $6.41\times10^{-17}$ | -8.72 | $5.53\times10^{-5}$ | 22.45 | $9.79\times10^{-3}$ | -0.09 | $7.55\times10^{-1}$ |
| 6 | 75.98 | $2.85\times10^{-1}$ | 7.08 | $4.97\times10^{-2}$ | 26.59 | $6.87\times10^{-2}$ | 0.63 | $1.87\times10^{-1}$ |
| 7 | 71.19 | $7.69\times10^{-3}$ | 6.45 | $4.20\times10^{-6}$ | -6.29 | $2.29\times10^{-1}$ | -0.18 | $2.85\times10^{-1}$ |
| 8 | 191.68 | $8.97\times10^{-6}$ | -1.24 | $5.89\times10^{-1}$ | -28.81 | $5.54\times10^{-4}$ | 0.29 | $2.81\times10^{-1}$ |

**Table S6: Fixed effects model terms for associations with Bayley Language scores**

| model terms | intercept | | age | | birth status | | Language | |
|---|---|---|---|---|---|---|---|---|
| state | β | p | β | p | β | p | β | p |
| 1 | 222.51 | $1.29\times10^{-8}$ | 2.20 | $3.27\times10^{-1}$ | -36.29 | $4.30\times10^{-5}$ | -0.67 | $3.02\times10^{-3}$ |
| 2 | 183.76 | $2.58\times10^{-4}$ | 3.96 | $1.76\times10^{-1}$ | 22.22 | $4.46\times10^{-2}$ | 0.36 | $2.08\times10^{-1}$ |
| 3 | 202.84 | $2.58\times10^{-4}$ | -1.55 | $6.30\times10^{-1}$ | -25.22 | $3.86\times10^{-2}$ | 0.72 | $1.96\times10^{-2}$ |
| 4 | 502.20 | $4.08\times10^{-20}$ | -9.52 | $2.83\times10^{-3}$ | 35.89 | $2.74\times10^{-3}$ | -0.85 | $5.10\times10^{-3}$ |
| 5 | 342.36 | $2.66\times10^{-17}$ | -9.28 | $4.78\times10^{-5}$ | 31.04 | $9.68\times10^{-4}$ | 0.19 | $4.40\times10^{-1}$ |
| 6 | 70.37 | $2.80\times10^{-1}$ | 8.07 | $2.85\times10^{-2}$ | 13.10 | $3.87\times10^{-1}$ | 0.41 | $2.87\times10^{-1}$ |
| 7 | 81.39 | $9.84\times10^{-4}$ | 6.66 | $2.59\times10^{-6}$ | -11.93 | $3.08\times10^{-2}$ | -0.35 | $1.30\times10^{-2}$ |
| 8 | 193.93 | $2.87\times10^{-6}$ | -0.66 | $7.84\times10^{-1}$ | -28.89 | $1.72\times10^{-3}$ | 0.21 | $3.73\times10^{-1}$ |
